# The Nuclear and Mitochondrial Genomes of *Amoebophrya* sp. ex *Karlodinium veneficum*

**DOI:** 10.1101/2024.09.05.611473

**Authors:** Wesley DeMontigny, Tsvetan Bachvaroff

**Affiliations:** Department of Cell Biology and Molecular Genetics, University of Maryland, College Park, MD 20742, USA; Institute for Marine and Environmental Technology, University of Maryland Center for Environmental Sciences, Baltimore, MD 21202, USA

**Keywords:** *Amoebophrya*, syndineans, Marine Alveolates, nonstop genetic code, UGA, dinoflagellate, intracellular parasite

## Abstract

Dinoflagellates are a diverse group of microplankton that include free-living, symbiotic, and parasitic species. *Amoebophrya*, a basal lineage of parasitic dinoflagellates, infects a variety of marine microorganisms, including harmful-bloom-forming algae. Although there are currently three published *Amoebophrya* genomes, this genus has considerable genomic diversity. We add to the growing genomic data for *Amoebophrya* with an annotated genome assembly for *Amoebophrya* sp. ex *Karlodinium veneficum*. This species appears to translate all three canonical stop codons contextually. Stop codons are present in the open reading frames of about half of the predicted gene models, including genes essential for cellular function. The in-frame stop codons are likely translated by suppressor tRNAs that were identified in the assembly. We also assembled the mitochondrial genome, which has remained elusive in the previous *Amoebophrya* genome assemblies. The mitochondrial genome assembly consists of many fragments with high sequence identity in the genes but low sequence identity in intergenic regions. Nuclear and mitochondrially-encoded proteins indicate that *Amoebophrya* sp. ex *K. veneficum* does not have a bipartite electron transport chain, unlike previously analyzed *Amoebophrya* species. This study highlights the importance of analyzing multiple genomes from highly diverse genera such as *Amoebophrya*.

**Summary:** This new long-read assembly demonstrates the remarkable diversity found within *Amoebophrya*. Despite being assigned the rank of genus, the available genome assemblies indicate significant variation in gene content, AT content, genetic codes, and potentially mitochondrial biology. Furthermore, this study contributes to the expanding list of organisms that contextually translate all three canonical stop codons. Although the mechanisms underlying such a genetic code remain elusive, the relative ease of culturing *Amoebophrya* suggests it may be useful as a model organism for future research on this subject.

## Introduction

Dinoflagellates (Alveolata, Myzozoa) are ecologically important microplankton found in most aquatic environments. The core dinoflagellates (*sensu* Janouskovec et al., 2016) are predominantly free-living or symbiotic, including common phytoplankton, coral symbionts (Freudenthal, 1962; Blank and Trench, 1986), and voracious microbial predators (Gaines and Taylor, 1984; Jacobson and Anderson, 1986). However, the base of the dinoflagellate tree is teeming with parasitic species; these parasites are predominantly known from environmental amplicon sequencing (Groisillier et al., 2008; Guillou et al., 2008) and collectively infect metazoans, ciliates, rhizarians, and other dinoflagellates (Cachon and Cachon, 1987).

*Amoebophrya*, a parasitic lineage in the MAGII/MALVII rDNA sequence clade (Guillou et al., 2008), is ubiquitous in marine environments and infects various dinoflagellate hosts (Cachon and Cachon, 1987). The life cycle begins as a biflagellate infective dinospore, which adheres to the surface of its host and penetrates its membrane (Cachon and Cachon, 1987; Miller et al., 2012). The parasite travels into the host nucleus or cytoplasm and develops into a multinucleate, beehive-like structure as it feeds on the cell (Cachon and Cachon, 1987; Fritz and Nass, 1992; Miller et al., 2012). Finally, the infection culminates when the host is violently lysed by plasmodial, multiflagellate, vermiform *Amoebophrya*, which subsequently swims off as it undergoes cytokinesis to produce new dinospores (Cachon and Cachon, 1987; Miller et al., 2012).

*Amoebophrya* is particularly noteworthy in its ability to infect dinoflagellates that cause harmful algal blooms (Coats et al., 1996; Coats et al., 1999; Park et al., 2004; Chambouvet et al., 2008; Chambouvet et al., 2011; Place et al., 2012; Velo-Suárez et al., 2013). For example, *Amoebophrya* sp. ex *Karlodinium veneficum* (*Amoebophrya* sp. Kv), the target of this sequencing study, is a species that infects *Karlodinium veneficum*, a small ∼12 micron, photosynthetic alga that feeds on dinoflagellates, cryptophytes, and animals (Place et al., 2012; Yang et al., 2020). *Karlodinium veneficum* secretes ichthyotoxic, hemolytic, and cytotoxic compounds known as karlotoxins (Deeds et al., 2002; Deeds et al., 2006), which decrease grazing pressure from zooplankton and aid in the capture of *K. veneficum*’s prey (Adolf et al., 2007). During a bloom of *K. veneficum*, karlotoxin secretion and rapid oxygen consumption by the algal mass can potentially kill large amounts of fish, with the largest fish kill attributed to *K. veneficum* being 30,000-50,000 fish (Place et al., 2012). *Amoebophrya* sp. Kv evades these toxins (Place et al., 2006) and is thought to aid in the termination of the bloom (Place et al., 2012).

Three *Amoebophrya* genomes are currently publicly available (*Amoebophrya* sp. A25, A120, and AT5) (John et al., 2019; Farhat et al., 2021), all from core dinoflagellate hosts. These genomes are strikingly diverse for organisms of the same genus. They contain a large number of species-specific genes, and even closely related species like A25 and A120 share less than half of their predicted proteins (Farhat et al., 2021). The genomes of some species have canonical introns (John et al., 2019), while others have many non-canonical splice sites and invasive intronic elements (Farhat et al., 2021).

Although there is no publicly available genome assembly for *Amoebophrya* sp. Kv, Bachvaroff (2019) analyzed a short-read-based, fragmented genomic survey and transcriptome. This genome differed from the previously published ones in two significant ways. First, about half of the annotated, transcribed genes in *Amoebophrya* sp. Kv were seemingly interrupted by all three canonical stop codons (UAA, UAG, and UGA), including genes of large and well-studied protein families like dynein heavy chains (Bachvaroff, 2019). The stop codons were not edited out of the stop-codon-containing genes (SCG) transcripts, and tRNAs recognizing UAA and UAG codons were identified (Bachvaroff, 2019). These traits suggested that, unlike the other *Amoebophrya* species, *Amoebophrya* sp. Kv uses an ambiguous genetic code - where canonical stop codons can act as sense codons depending on their context. Second, this assembly included the mitochondrial genome of *Amoebophrya* sp. Kv (Bachvaroff, 2019), which has remained elusive in the other genome assemblies (Kayal and Smith, 2021); it has even been tentatively suggested that *Amoebophrya* lacks a mitochondrial genome altogether and that complex III of the electron transport chain is missing, splitting the electron transport chain into two (John et al., 2019). However, this does not seem to be true for *Amoebophrya* sp. Kv, which has cytochrome b encoded in its mitochondrial genome (Bachvaroff, 2019).

The original culture used by Bachvaroff (2019) has since been lost. The isolation of a new *Amoebophrya* strain and the development of Oxford Nanopore sequencing offer the opportunity to analyze a high-quality, independent genome of *Amoebophrya* sp. Kv. Therefore, we sought to validate the previous observations about *Amoebophrya* sp. Kv and add to the growing genomic data on *Amoebophrya*. Here, we present a draft genome of *Amoebophrya* sp. Kv, further evaluate the evidence for ambiguous genetic stop codons in this species and provide the most complete *Amoebophrya* mitochondrial contigs to date.

## Materials and Methods

### Culturing

The *Amoebophrya* strain used in this study was isolated from the Inner Harbor, Baltimore, in June of 2011, where a *K. veneficum* bloom was observed. Vibrant green-blue fluorescence from within the *K. veneficum* cells was observed using fluorescence microscopy, indicating infection by *Amoebophrya*. Water samples were filtered across an 8 μm Millipore membrane, diluted, and added to naive cultured *K. veneficum* CCMP1975 in a 48-well plate. The naive host culture was grown at 20 °C with 100 μMoles Photons m^-2^s^-2^ and a 14h light/10h dark cycle in F/2 media (without silica) at a salinity of 15 ppt made from locally collected estuarine water. After one week, several wells contained green fluorescent hosts and visible fluorescent dinospores. Individual infected cells were picked with a drawn glass pipette, washed in filtered media, and added to new naive hosts in 48 well plates. A single clonal culture from a washed infected host cell was established and maintained on *K. veneficum*.

### DNA Sequencing and Assembly

Weekly culture changes with a 10:1 ratio of parasite spore to host cells (100,000 cells mL^-1^ versus 10,000 cells mL^-1^) of the host were diluted in fresh media to concentrations of 50,000 cells mL^-1^ of *Amoebophrya* spores and 5,000 cells mL^-1^ of naive host cells. Cell concentrations were measured using a Coulter counter with a 20 μm orifice. Week-old cultures at 100,000 spore cells mL^-1^ in a total volume of 150 mL were centrifuged at 10,000 g for 20 minutes, and the cell pellets were used for DNA isolation. Cells were lysed with a 2% CTAB detergent solution and incubated for 20 minutes at 50 °C. DNA was extracted from the crude lysate with two rounds of chloroform extraction and precipitation with two volumes of ethanol. The Short Read eliminator kit from PacBio was also used to increase average fragment size.

Each sequencing library used 1 μg of starting DNA with polishing of the DNA using the NEB Nanopore Sequencing Companion Kit, followed by ligation and size selection using the Nanopore Ligation Sequencing Kit V14. Sequencing was performed on a MinION sequencer on an R10 chip with multiple library loadings when the read output declined. Basecalling was performed using the Dorado pipeline’s “Super Accurate” model with duplex basecalling using the stereo model of predicted duplex reads on the ada GPU cluster at the University of Maryland Baltimore County. The genome was assembled using Canu 2.2 (Koren et al., 2017) with an expected genome size of 300 Mb and read parameters appropriate for an R10 Nanopore (*chp ’corMhapOptions=--threshold 0*.*8 --ordered-sketch-size 1000 --ordered-kmer-size 14’ correctedErrorRate=0*.*105*). ABySS-fac (Jackman et al., 2017) was used to generate assembly statistics.

Contigs were flagged as bacterial contaminants using MEGAN (Huson et al., 2007). The *Amobeophrya* subset of the metagenome assembly was determined by selecting non-bacterial contigs with at least 20x genomic coverage and 50% coverage from the *Amoebophrya* sp. Kv transcriptome and an AT proportion of 63-67%. The total metagenome assembly was visualized using the ggplot2 package (Wickam, 2016). The Bachvaroff (2019) *Amoebophrya* sp. Kv assembly was mapped to the new assembly using minimap2 (Li, 2018). The number of contigs from the old assembly that were at least 90% represented was counted using a custom script.

### Genome Annotation

Repeat libraries for the parasite contigs were generated with RepeatModeler (Smit and Hubley, 2008-2015) and masked with RepeatMasker (Smit et al., 1996-2010). Transfer RNA genes were identified using tRNAscan-SE with maximum sensitivity (Lowe and Eddy, 1997). The tRNAscan-SE predicted secondary structure of TrpCCA tRNAs was manually investigated for variation in the anticodon stem length. RNA-seq validated introns with a minimum anchor of 30 and minimum read coverage of 3 were extracted from the parasite contigs using Minimap2 (Kim et al., 2019) and Regtools (Cotto et al., 2023). The donor and acceptor splice site frequencies were visualized using WebLogo (Crooks et al., 2004).

The MAKER annotation pipeline (Cantarel et al., 2008) was provided with *de novo* assembled Trinity transcripts from Bachvaroff et al. (2014), repeats from RepeatMasker, proteins from other species of *Amoebophrya*, and a variety of myzozoan proteomes previously used to assemble gene families for *Amoebophrya* sp. A120 (Farhat et al., 2021; https://www.ncbi.nlm.nih.gov/datasets/genome/GCA_905178155.1/) and A25 (Farhat et al., 2021; https://www.ncbi.nlm.nih.gov/datasets/genome/GCA_905178165.1/), *Perkinsus marinus* (Unpublished; https://protists.ensembl.org/Perkinsus_marinus_atcc_50983_gca_000006405/), *Symbiodinium microadriaticum* (Aranda et al., 2016; http://smic.reefgenomics.org/), *Fugacium kawagutii* (Lin et al., 2015; http://web.malab.cn/symka_new/), *Breviolum minutum* (Shoguchi et al., 2013; http://marinegenomics.oist.jp/symb/viewer/info?project_id=21), *Plasmodium falciparum* strain 3D7 (Aurrecoechea et al., 2009; http://phlasmodb.org/plasmo/), *Toxoplasma gondii* strain ME49 (Kissinger et al., 2003; http://toxodb.org/), *Chromera velia* strain CCMP2878 (Woo et al., 20015; https://cryptodb.org/), *Vitrella brassicaformis* strain CCMP 3155 (Woo et al., 2015; https://cryptodb.org/), *Theileria equi* (Kappmeyer et al., 2012; https://piroplasmadb.org/), and *Cryptosporidium parvum* (Abrahamsen et al., 2004; https://cryptodb.org/).

Since MAKER fails to detect SCGs because it terminates gene models at the first in frame stop codon, a complementary approach was used to find those genes. Intergenic regions were extracted from MAKER and blasted against *Amoebophrya* sp. A120 and A25 proteomes with a minimum e-value of 1e-5. Genomic regions with hits were extracted and aligned with the proteins using MiniProt (Li, 2023). Only alignments longer than 100 amino acids were kept. Protein alignments lacking in-frame stop codons or frameshifts were fed into MAKER for another round of gene prediction. Regions lacking MiniProt alignments and MAKER-predicted genes were again aligned to the *Amoebophrya* proteomes for protein alignments. Finally, protein alignments lacking in-frame stop codons or frameshifts were fed into MAKER for a final round of gene prediction. Stop-codon-containing alignments were extracted from the MiniProt outputs. When protein alignments from different queries overlapped, only the best-scoring alignment was kept. Any alignments containing frameshifts were discarded from further analyses. The number of alignments supported by transcript evidence was determined by blasting the alignments to the Trinity transcriptome with a minimum percent identity of 90%; only alignments with 80% coverage by the transcriptome were considered supported.

In-frame stop codons of the alignments were masked, and the MAKER predictions and alignments were annotated by InterProScan (Jones et al., 2014) and assigned KEGG orthology using BlastKOALA (Kanehisa et al., 2016). Completeness of the gene predictions was assessed with BUSCO using the alveolata_odb10 database (Simao et al., 2015). OrthoFinder (Emms and Kelly, 2019) was used to find orthologous groups between the proteomes previously used in the MAKER annotation pipeline and the MAKER-predicted proteins and stop-codon-masked protein alignments. Proteins important for cellular processes were identified based on shared sequence identity, protein domains, and KEGG orthology. Select proteins and in-frame stop codons were visualized using the gggenome R package (Hackl et al., 2023).

### Mitochondrial Analysis

The *Hematodinium* mitochondrial proteins COXI, COXIII, and CYTB (CCE53570.1, CCE53572.1, CCE53571.1) were queried against the assembly using tBLASTn using the protozoan mitochondrial code. The best hits were labeled as putative mitochondrial proteins and scanned for functional domains using InterProScan (Jones et al., 2014). These proteins were blasted back to the assembly, and contigs containing hits with a minimum e-value of 1e-75 and 95% shared identity were labeled as putative mitochondrial contigs. Mitochondrial contigs were searched for additional protein-encoding genes using InterProScan (Jones et al., 2014). Inverted repeats were identified using the repeat-match program packaged with MUMmer (Marçais et al., 2018). Nuclear-encoded proteins belonging to the mitochondrial respiratory chain were searched for in the MAKER-predicted gene models and unincorporated protein alignments. Select mitochondrial contigs were visualized using the gggenome R package (Hackl et al., 2023).

## Results

### DNA Extraction and Assembly

The input for the assembly was 5.3 million Nanopore reads with an N50 of 9 kb and a total length of 27.7 Gb. The overall assembly of *Amoebophrya*, host, and bacterial contigs using Canu was 344.4 Mb in 3018 contigs with an N50 of 3.41 Mb and an L50 of 32 (Figure 1). The cultured metagenomic assembly included three distinct bins of data. A total of ∼121 Mb in 39 contigs were attributed to co-cultured bacteria based on SSU ribosomal RNA strong BLAST hits to the 16S_microbial database from NCBI. The *Amoebophrya* subset of the metagenome assembly consisted of 48 contigs with a total length of 127 Mbp and N50 of 4.1 Mb (Table 1).

**Figure 1.**
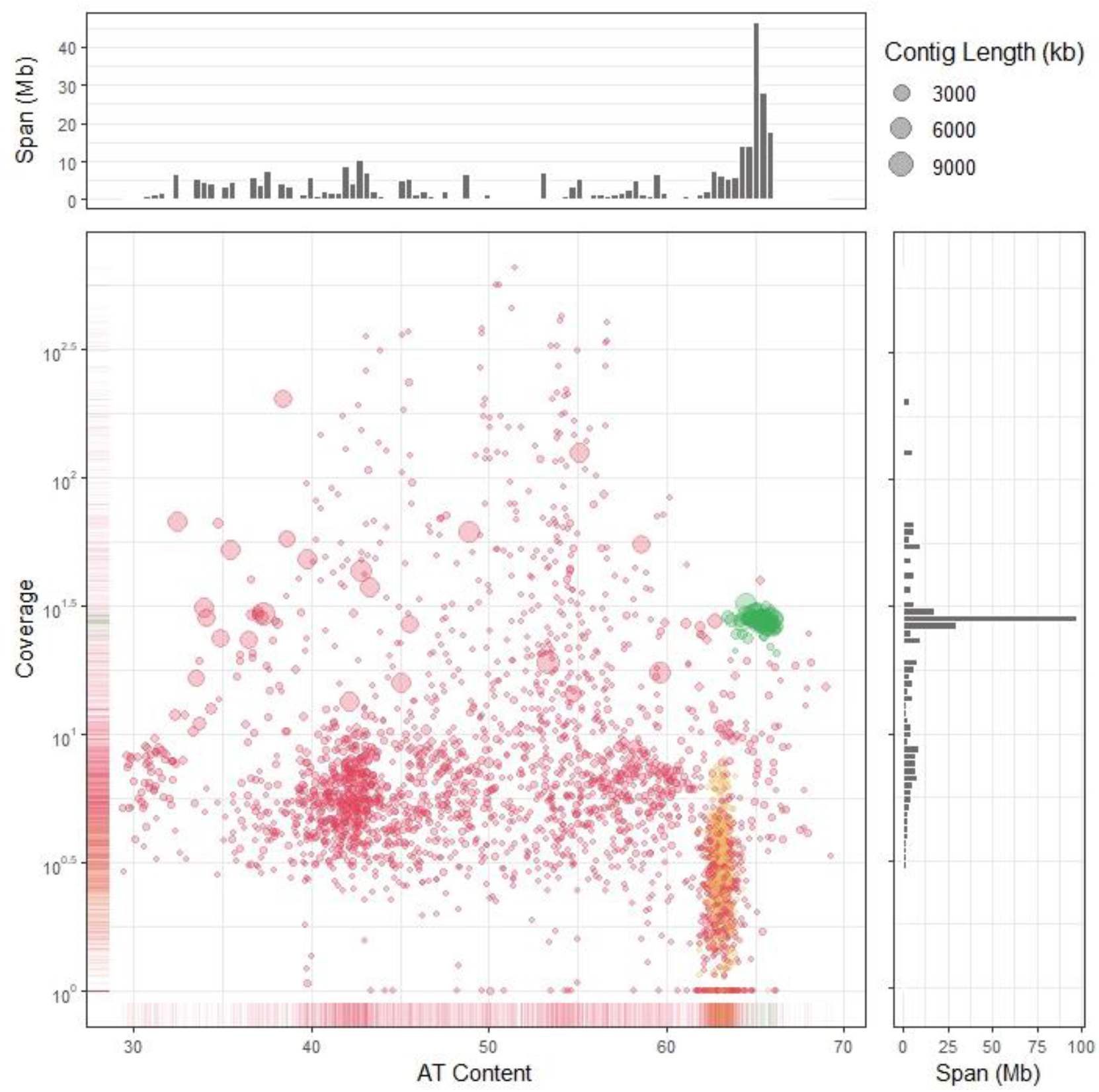
A plot of the base coverage against the AT content of the contigs in the Canu metagenome assembly. The side plots indicate the number of megabases represented in a region of the plot. The genomic and mitochondrial contigs of *Amoebophrya* sp. ex *K. veneficum* are colored green and orange, respectively, and bacterial and *Karlodinium veneficum* contigs are colored red.

**Table 1.** Comparisons of the assemblies and annotation of the *Amoebophrya* sp. ex *Karlodinium veneficum* and *Amoebophrya* sp. A120, A25, and AT5.

|  | **Kv** | **A120** | **A25** | **AT5** |
| --- | --- | --- | --- | --- |
| **Assembly** |  |  |  |  |
| Number of scaffolds/contigs | 48 | 50 | 556 | 2351 |
| Cumulative size (Mb) | 127 | 115.5 | 116 | 87.7 |
| Scaffold/Contig N50 | 4.119 Mb | 9.243 Mb | 1.082 Mb | 83.9 kb |
| Scaffold/Contig max. size | 11.551 Mb | 16.512 Mb | 3.013 Mb | 1.914 Mb |
| AT Content | 65.2 | 48.8 | 52.2 | 54.5 |
| **Protein-Encoding Genes** |  |  |  |  |
| Number | 8,295 | 26,441 | 28,091 | 19,925 |
| Average gene length (bp) | 2,267 | 3,482 | 2,965 | 2,782 |
| Average protein length (aa) | 426 | 591 | 446 | 654 |

### Genome Annotation

Repeat masking using RepeatModeler and RepeatMasker masked 13.7 Mb (10.72%) of parasite contigs. No DNA transposons were identified, but 848 retroelements across 2.1 Mb were found. There were 2.0 Mb of LTR elements dispersed throughout the genome, but most repeats were unclassified. The 216 different unclassified repeats spanned 8.6 Mb (6.8%) of the genome and had a wide range of AT bias. Simple sequence repeats and low-complexity regions represented only 2.1 Mb of the assembled data.

The *Amoebophrya* sp. Kv contigs were scanned for tRNAs with maximum sensitivity using tRNAscan-SE. A total of 163 tRNAs and 24 pseudo-tRNAs were predicted. Four putative cognate suppressor tRNAs were found in the *Amoebophrya* sp.Kv assembly - two cognate tRNAs for UAA and UAG. In the assemblies of *Amoebophrya* sp. 120, A25, and AT5, only one pseudo-tRNA identified as a stop-suppressor was detected in A25. One of the seven predicted TrpCCA tRNAs had a 4 bp anticodon stem, and the bases typically forming the fifth pair of the anticodon stem are U26 and C42 (Supplemental Figure 1).

The Rfam website was used to identify four contigs with 6773 base rDNA repeats containing SSU, ITS1, 5.8S, ITS2, and LSU repeated from 6 to 16 times in a row. There were no credible sequence differences in the 45 complete or near complete SSU and LSU regions in the four contigs. Although tig00002270 starts with a variant SSU, read mapping does not support this result. After the LSU, there were consistently 5S RNA and SL sequences between the rDNA regions, followed by the next SSU region. Introns validated by RNA-seq data were predominantly GT/AG splicing (Figure 2) and rarely AT/AC, GC/AG, or AT/TC splicing. This observation is supported by the identification of U1, U2, U4, U5, and U6 snRNAs in the parasite assembly.

**Figure 2.**
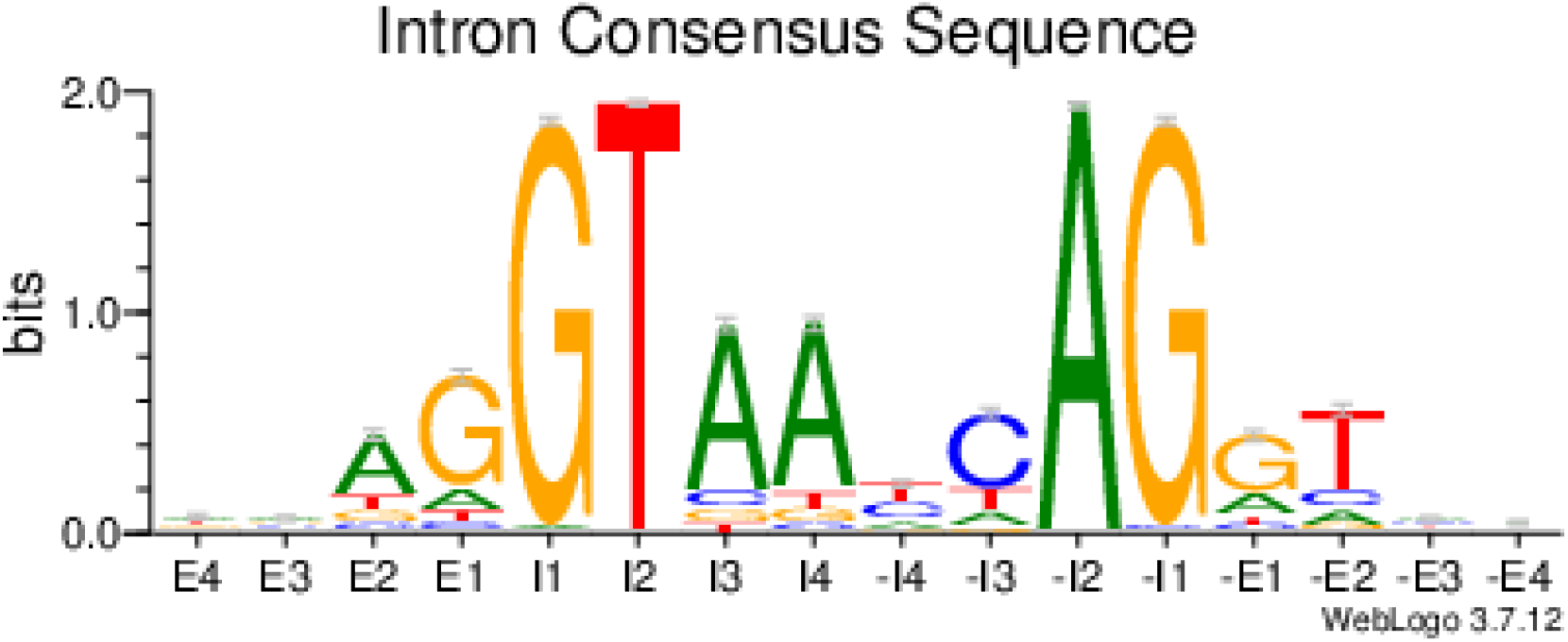
A logo plot indicating the sequence motifs at the end of exons (E4 - E1), the beginning of introns (I1 - I4), the end of introns (-I4 - -I1), and the beginning of exons (-E1 - -E4).

Two complementary approaches were used for the prediction of protein-coding genes. One was based on MAKER, which was restricted to using the standard genetic code, and a second method was based on MiniProt for predicting genes with apparent stop codons in their open reading frames (SCGs). Based on protein identity and transcriptome-based evidence, MAKER predicted 4,668 protein-encoding genes, and 2,367 proteins contained predicted Pfam domains. The average MAKER gene was 2,063 bp long, contained 2.6 exons, and encoded a 330.8 amino acid protein (Table 1). MiniProt revealed 3,627 SCGs, with 2,462 represented in the previously published transcriptome (Supplemental Material) and 3,544 having predicted Pfam domains. The average SCG was 549 amino acids long, and the average number of stops in each SCG was 13.6 (58% UAA, 32.3% UAG, 9.7% UGA). When SCGs were excluded, the annotation only had 59.7% BUSCO completeness; including the SCGs raised the BUSCO completeness score to 79.5%. For comparison, the alveolate BUSCO completeness of the predicted genes for AT5, A120, and A25 are 85.6%, 94.7%, and 90.6%, respectively. Of the 8,295 amino acid sequences provided to OrthoFinder, 6,837 were assigned to 4,798 orthogroups. Of the 4,798 orthogroups, 2,181 contained SCGs and did not include MAKER-predicted genes.

### Mitochondrial Analyses

The assembly contained a large number of high-quality tBLASTn hits for mitochondrially-encoded proteins from *Hematodinium* (CCE53570.1, CCE53572.1, CCE53571.1) corresponding to *coxI, coxIII*, and *cytB* that shared little identity with those of *K. veneficum* (AAQ04825.2, ABR15108.1, ABR15096.1). The best-scoring BLAST hits were selected as putative mitochondrial proteins for *Amoebophrya* sp. ex *K. veneficum*.

The mitochondrial proteins occur on 491 AT-rich contigs in the assembly (Figure 1).

However, the contigs are quite dissimilar; some only encode a single protein, while others encode all three (Figure 3). The mitochondrial contigs have relatively low base coverage (1x-8.05x), with lengths ranging from 3339 bp to 56,915 bp and little-to-no RNA-seq coverage (Supplemental Figure 2). When contigs were searched for inverted repeats with lengths greater than 50 nt, 3196 pairs of varying lengths (50-5332 bp) were detected; clustering with CD-HIT at 95% identity revealed 122 different clusters of inverted repeats.

**Figure 3.**
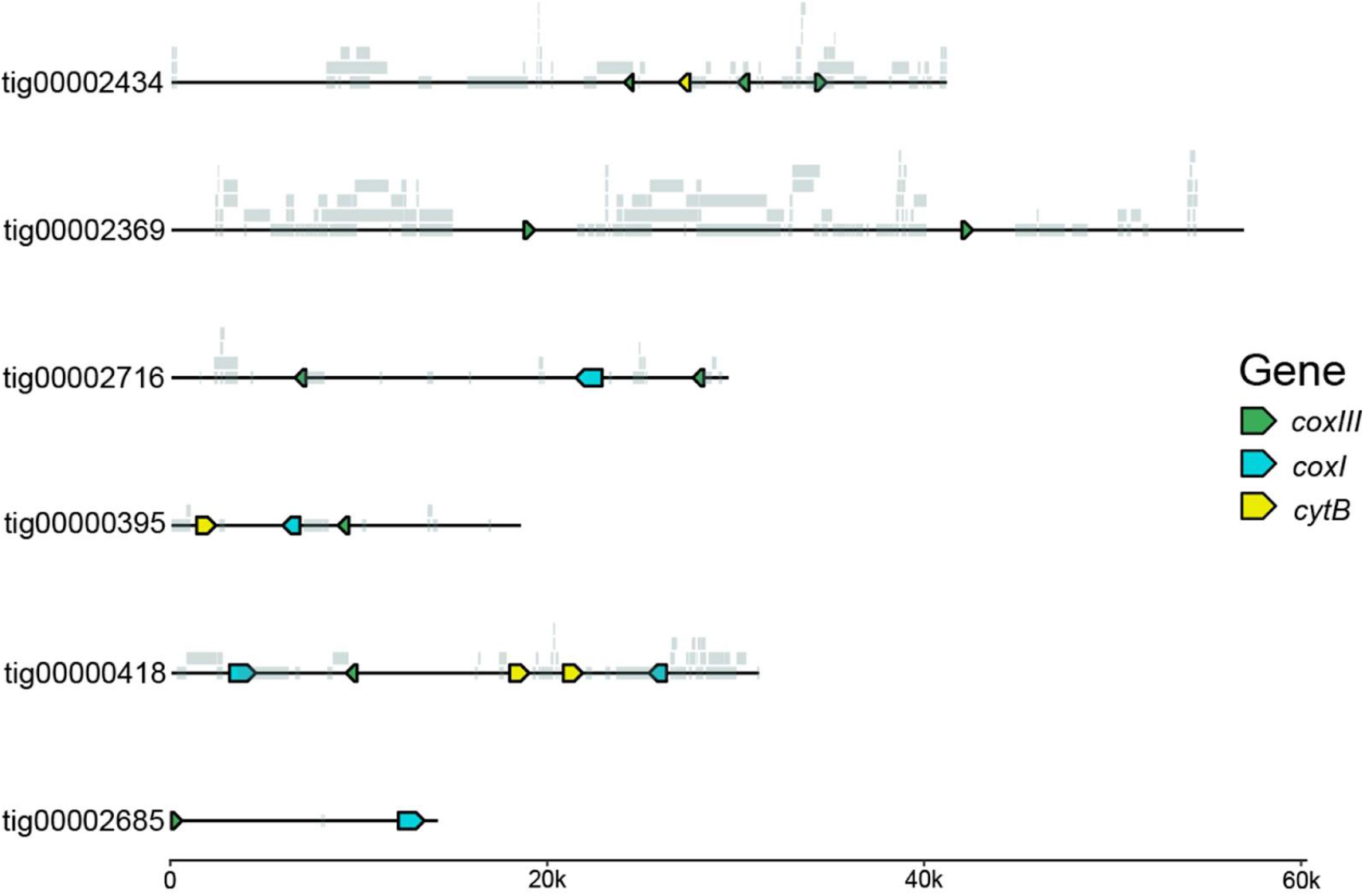
Selected mitochondrial contigs from *Amoebophrya* sp. ex *Karlodinium veneficum* with fragmented and full copies of *coxI, coxIII*, and *cytb*. Grey blocks along the contigs indicate the location of inverted repeats.

The nuclear genome contained proteins from all major mitochondrial electron transport chain complexes except Complex I. Complex II (succinate dehydrogenase) and the F1 subunit of Complex V (ATP synthase alpha, beta, gamma, delta, and delta OSCP subunits) were complete. Complex III (cytochrome c1, Rieske iron-sulfur protein, mitochondrial processing-peptidase subunits alpha and beta, and cytochrome b-c1 complex subunit 7) and Complex IV (cytochrome c oxidase subunit 2, 5b, 6b, 11, 15, and 19) were only partially represented, and the F0 subunit of Complex V was undetected. Genes encoding cytochrome c, alternative NADH dehydrogenase, and alternative oxidase were also found. All the nuclear encoded proteins lacked in-frame stop codons except for the alternative NADH dehydrogenase and cytochrome c oxidase subunits 11, 15, and 19.

## Discussion

### The Nuclear Genome of *Amoebophrya* sp. ex *Karlodinium veneficum*

In eukaryotes, canonical stop codons typically terminate translation when the eRF1 protein recognizes the stop codon and releases the nascent peptide chain (Jackson et al., 2012). However, several eukaryotes have reassigned canonical stop codons to code for amino acids by using suppressor tRNAs that pair with stop codons. Such codon reassignment has been reported in diverse lineages, including mitochondria (Barrel et al., 1979), viruses (Borges et al., 2022), bacteria (Inamine, 1990), green algae (Schneider et al., 1989; Schneider and de Groot, 1991), diplomonads (Keeling and Doolittle, 1996), and ciliates (Hoffman et al., 1995; Lozupone et al., 2001). However, some species use ambiguous stop codons recognized by both suppressor tRNAs and eRF1, allowing the codon to code for an amino acid or terminate translation depending on the context. Organisms that utilize ambiguous stop codons include the trypanosomatid genus *Blastocrithidia* (Kachale et al., 2023), the heterotrich ciliate *Condylostoma magnum* (Swart et al., 2016), and karyorelict ciliates (Swart et al., 2016; Seah et al., 2022). Although a mechanism has been proposed for how certain ciliates define stop/coding contexts of ambiguous stop codons (Swart et al., 2016), it seems unlikely that this is conserved across the distantly related lineages that utilize this sort of genetic code.

Here, we present the genome assembly of a novel strain of *Amoebophrya* sp. Kv, which is the second most contiguous *Amoebophrya* genome assembly currently available (Table 1). This assembly is also a significant improvement on the prior assembly by Bachvaroff (2019), condensing 8,801 contigs into just 48. The genome of *Amoebophrya* sp. Kv is much more AT-rich than the other published *Amoebophrya* genomes (Table 1), and *Amoebophrya* sp. Kv primarily uses canonical GT/AG splicing patterns (Figure 2). This splicing pattern is consistent with what has been reported for *Amoebophrya* sp. AT5 (John et al., 2019) and further supports the strange intronic elements of A120 and A25 being confined to only a subset of the genus (Farhat et al., 2021).

Similar to the 2019 genomic survey, this assembly contained cognate tRNAs for UAA and UAG codons. Although we detected no UGA cognate tRNA, *Blastocrithidia nonstop*, a trypanosomatid that has repurposed all three canonical stop codons, also has no dedicated UGA cognate tRNA in its genome. Instead, it can translate UGA using a Trp_CCA_ tRNA with a 4 bp anticodon stem (AS) rather than the typical 5 bp AS (Kachale et al., 2023). The suppression of UGA by a 4 bp AS Trp_CCA_ tRNAs was experimentally transferable to *Trypanosoma brucei, Condylostoma magnum*, and *Saccharomyces cerevisiae*, suggesting that this is a characteristic of these tRNAs in general (Kachale et al., 2023). Moreover, such 4 bp AS suppressor Trp_CCA_ tRNAs appear to be present in *Blepharisma* and *Loxodes* (Swart et al., 2024), and precedent for 4 bp AS suppressor tRNAs in *Escherichia coli* dates back to 1994 (Shultz and Yarus, 1994a; Shultz and Yarus, 1994b; Swart et al., 2024). Like these organisms, *Amoebophrya* sp. Kv also has a Trp_CCA_ tRNA with a 4 bp AS; interestingly, the two bases that would otherwise form the fifth pair of the anticodon stem are identical to those of the tRNA that suppressed UGA in *C. magnum* (Kachale et al., 2023). We, therefore, feel that this Trp_CCA_ tRNA is likely the mechanism for UGA suppression.

*Amoebophrya* sp. Kv possesses two eRF1 homologs that are identical to other *Amoebophrya* species at amino acid motifs associated with stop codon recognition or polypeptide release. However, in all *Amoebophrya* species analyzed, the conserved NIKS motif has been modified to RIKS. Mutations at this asparagine residue are known to modify the termination efficiency of UAA and UAG (Frolova et al., 2002), but the exact effect of this mutation is impossible to assess from sequence data alone. Furthermore, modification of such a highly conserved residue in *Amoebophrya* with a canonical genetic code indicates that it does not prevent termination at UAA and UAG. Combined with the previously mentioned suppressor tRNAs, this suggests that all three canonical stop codons are recognized by both tRNAs and eRF1 in *Amoebophrya* sp. Kv.

Nearly half of the predicted proteins in this assembly contained in-frame stop codons, which is consistent with the inference that *Amoebophrya* sp. Kv utilizes ambiguous stop codons. These SCGs are a genuine characteristic of this species, evidenced by their presence in genomes of different *Amoebophrya* sp. Kv isolates that were sequenced using different technologies. Furthermore, the SCGs are unlikely to be pseudogenes. This is supported by the observation that 2,181 orthogroups contained an SCG but no MAKER-predicted proteins. If the SCGs in these orthogroups are pseudogenes, it would mean that *Amoebophrya* sp. Kv performs these orthogroups’ functions via alternative mechanisms or has lost the need for that function altogether. Parasite genomes are often characterized by a loss of function and increased dependence on the host, and many orthogroups may be functionally redundant to some extent. However, several SCGs are critical for cellular processes that cannot be easily delegated to the host and are not known to be functionally redundant. This is especially true for proteins involved in DNA replication (e.g., RNase H, Top1-3, and the catalytic subunits of DNA polymerases alpha and delta), DNA repair (e.g., 8-oxoguanine DNA glycosylase, DNA polymerase kappa, and RAD54), and pre-mRNA splicing (e.g., splicing factor 3B subunits 1, 3, and 4) (Figure 4). If these SCGs are pseudogenes, this would imply a substantial loss of function; *Amoebophrya* sp. Kv would presumably be hindered in its ability to replicate DNA, repair common DNA lesions, and splice introns. The improbability of these scenarios suggests that the SCGs are indeed translated into proteins. Ultimately, the translation of all three stop codons is the most parsimonious explanation for our observations. From the isoforms of the suppressor tRNAs and previous codon usage analyses (Bachvaroff, 2019), it seems likely that UGA codes for tryptophan and UAA/UAG codes for glutamine, but this will need to be confirmed experimentally. Translation termination at one or more of these codons is likely contextual, but determining which codon(s) signal for termination and the required context will require additional study.

**Figure 4.**
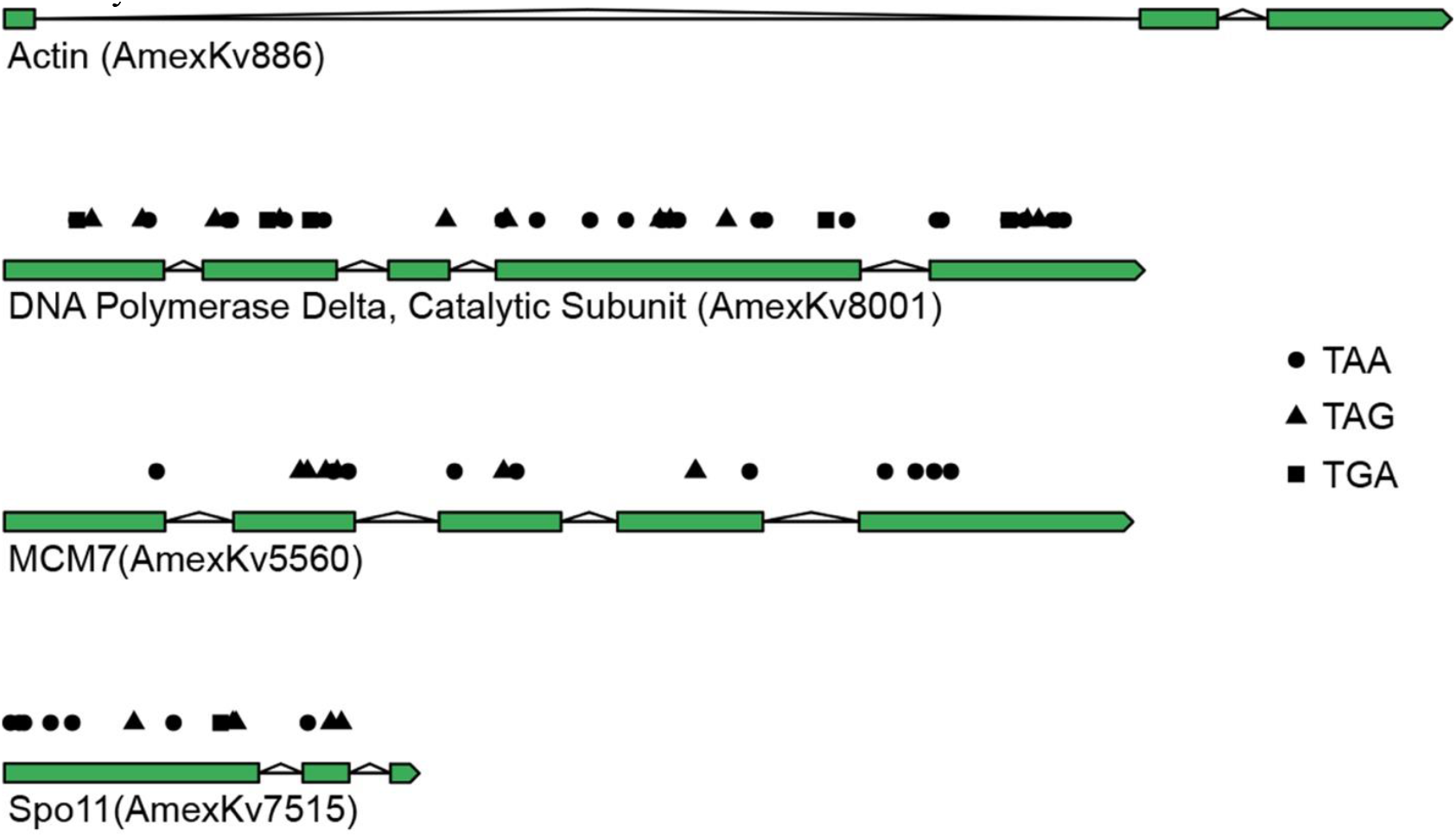
Select gene models from *Amoebophrya* sp. ex *Karlodinium veneficum*. Here, actin is a gene model that lacks in-frame stop codons. Stop codons present in the open reading frame are indicated by circles (TAA), triangles (TAG), and squares (TGA) above the gene model.

Ambiguous stop codons make it challenging to discriminate between translated and untranslated reading frames; this makes gene calling particularly difficult, especially in a lineage with many species-specific genes. It is likely that there are more protein-encoding genes than the 8,295 we predicted. However, because these unidentified genes lack strong sequence identity with known myzozoan genes and likely contain many stop codons, we were unable to find an efficient way to identify them. Furthermore, our gene predictions may contain some pseudogenes, missed introns, fragmented gene models, and other artifacts of high throughput annotations. Refining our gene models requires knowledge of what constitutes a true stop codon, which is difficult to determine with the available data.

### The Mitochondria of *Amoebophrya* sp. ex. *Karlodinium veneficum*

Dinoflagellates and their sister, the apicomplexans, are notable for having some of the most gene-poor mitochondrial genomes on the eukaryotic tree (Gagat et al., 2017), encoding for only three proteins (COXI, COXIII, and CYTB) and scattered fragments of rDNA (Gagat et al., 2017). The mitochondrial genome of *Amoebophrya* has not been detected in the AT5, A120, or A25 genome assemblies, except for a short fragment resembling *coxI* in A120 and AT5 (Kayal and Smith, 2021). This caused a debate about whether *Amoebophrya* lacked a mitochondrial genome altogether (Kayal and Smith, 2021; John et al., 2019). Although only mentioned in passing, the mitochondrial genome and base editing of *Amoebophrya* sp. ex *Karlodinium veneficum* has been previously reported (Bachvaroff, 2019). The controversy surrounding *Amoebophrya*’s mitochondrial genome prompted further examination here.

The present study detected nearly 500 contigs containing genes for *Amoebophrya* sp. Kv’s *coxI, coxIII*, and *cytb* (Supplemental Figure 2 and Figure 3). Although we lack localization data for these sequences and recognize that these contigs could be from the nuclear genome, we strongly favor their assignment to the mitochondrial genome. The gene content of these contigs is typical for a myzozoan mitochondrial genome, and despite the large size of some contigs, they lack any sign of nuclear-encoded genes.

The protein-encoding genes in each mitochondrial contig had highly conserved sequence identity, but outside of the genes, the contigs varied greatly. All three protein-encoding genes were present in some mitochondrial contigs, while others contained only a subset of the genes or fragmented copies. Additionally, the intergenic regions often consisted of inverted repeats, the exact sequence of which varied between 122 repeat families. The extreme variation in mitochondrial contigs raises the possibility that a heterogeneous population of mitochondrial fragments contributes to the functionality of a single mitochondrion in *Amoebophrya* sp. Kv. Alternatively, the mitochondrial genome of *Amoebophrya* sp. Kv may differ significantly from individual to individual and exist only at low copy numbers; this would make it difficult to assemble a complete mitochondrial genome from many different dinospores. However, given that the sequenced cultures descend from a single infected host and are passed through a population bottleneck with each transfer, it would be surprising to observe such a diversity of mitochondrial genomes among individuals. Ultimately, more work is needed to assess if *Amoebophrya*’s mitochondrial genome is fragmented and what the relationship is among these multiple contigs.

Mitochondrial proteins from all electron transport chain (ETC) complexes besides complex I were encoded in the nuclear genome. All complex III respiratory and core subunits were identified, which are notably missing in the gene predictions of *Amoebophrya* sp. AT5, A120, and A25. The absence of complex III proteins in these genomes led to the suggestion that *Amoebophrya* has a bipartite ETC (John et al., 2019). While the ETC may indeed be split in some *Amoebophrya* species, this does not appear to be the case for *Amoebophrya* sp. Kv is, therefore, not a general property of the genus. Further investigations are required to determine if *Amoebophrya* has diverse mitochondrial biology or if it is more conventional than was previously thought.

## Data Availability

Analyses that are more complicated than singular commands are available as scripts and can be found at https://github.com/Wesley-DeMontigny/AmexKv-Genome-Scripts. Supplemental figures and data files can be found at doi 10.6084/m9.figshare.26940256. The NCBI bioproject accession is PRJNA1103200 and the raw reads can be found at SRR28830816-8 and SRR28782276-8.

## Acknowledgements

We would like to thank Shehre Banoo Malik, John Mattick, and Charles Francis Delwiche for their comments on the manuscript. WCD is currently a graduate student in CFD’s lab. The hardware used in the computational studies is part of the UMBC High Performance Computing Facility (HPCF). The facility is supported by the U.S. National Science Foundation through the MRI program (grant nos. CNS–0821258, CNS–1228778, OAC–1726023, and CNS–1920079) and the SCREMS program (grant no. DMS–0821311), with additional substantial support from the University of Maryland, Baltimore County (UMBC). See hpcf.umbc.edu (accessed on 7 July 2024) for more information on HPCF and the projects using its resources. The BioAnalytical Services Laboratory (BASLab) at the Institute of Marine and Environmental Technology (IMET) was used for Nanopore GridION and MinION long-read sequencing. See https://www.umces.edu/baslab for more information on the BASLab capabilities.

## Funding

Funding was provided by the IMET Angel Investors fund and the J. Unger Vetlessen foundation.

## Conflicts of Interest

The authors declare that they have no conflicts of interest.

## Supplemental Figures

**Supplemental Figure 1.**
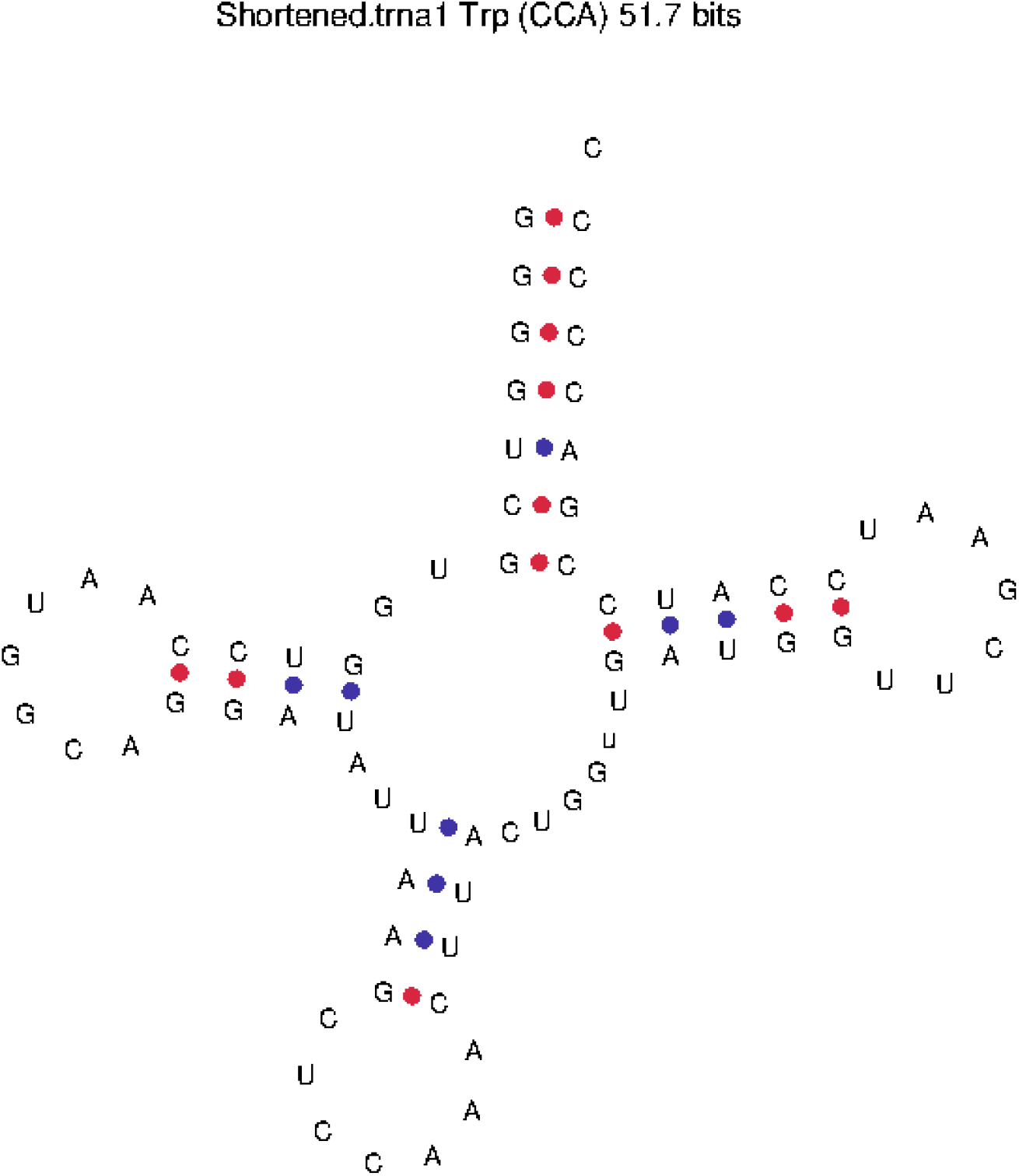
The tRNAscan-SE predicted structure for the Trp_CCA_ tRNA with a 4 bp anticodon stem encoded in the *Amoebophrya* sp. ex *Karlodinium veneficum* genome.

**Supplemental Figure 2.**
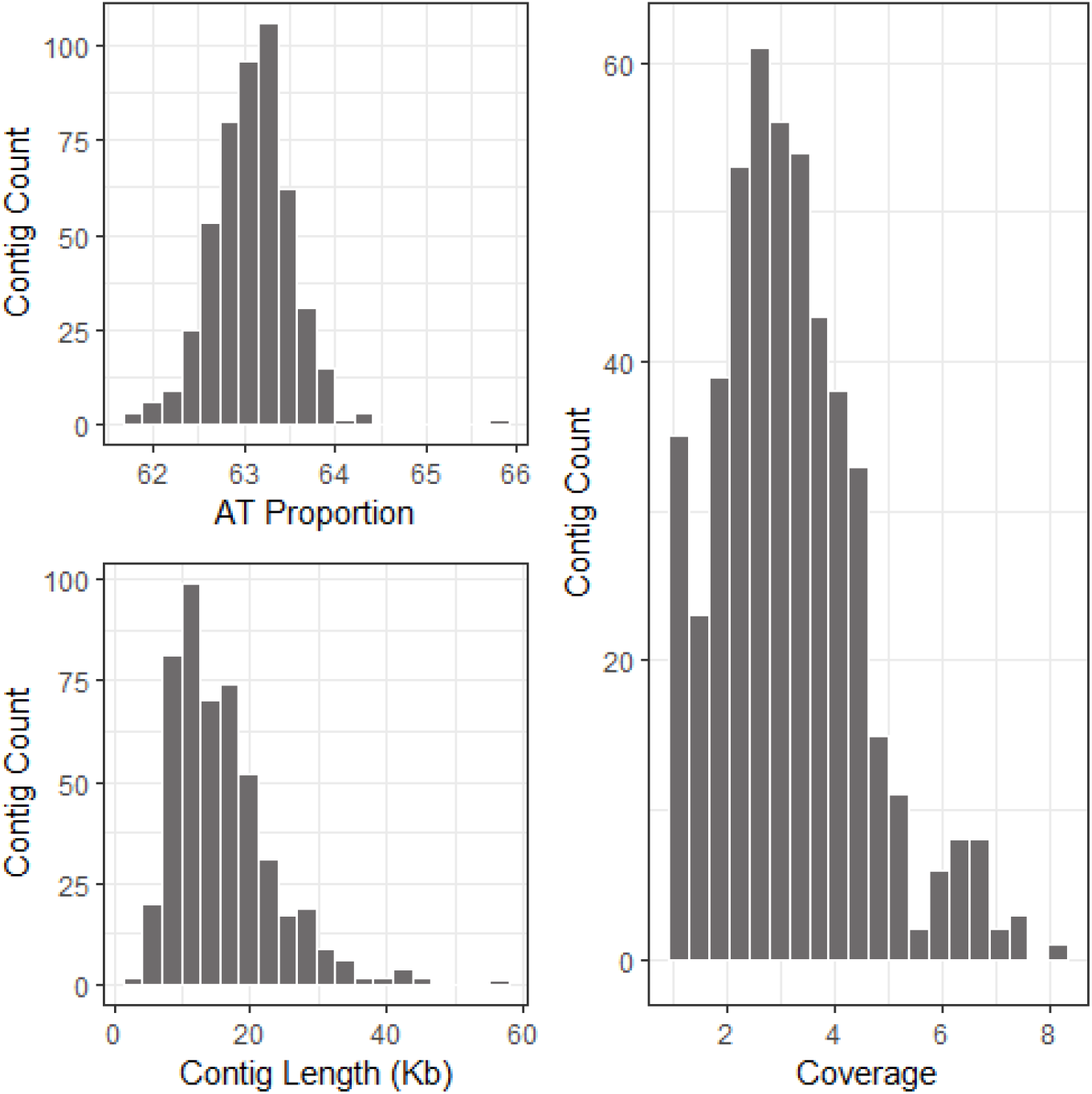
The distributions of AT proportion, length, and coverage of the putative mitochondrial contigs for *Amoebophrya* sp. ex *Karlodinium veneficum*.

